# The first observation of osmotically neutral sodium accumulation in the myocardial interstitium

**DOI:** 10.1101/2021.07.02.450864

**Authors:** I. Artyukov, G. Arutyunov, M. Bobrov, I. Bukreeva, A. Cedola, D. Dragunov, R. Feshchenko, M. Fratini, V. Mitrokhin, A. Sokolova, A. Vinogradov, A. Gianoncelli

## Abstract

The aim of this study was the detection and quantification of the Na+ depositions in the extracellular matrix of myocardial tissue, which are suggested to be bound by negatively charged glycosaminoglycan (GAG) structures. The presented experimental results are based on high resolution X-ray fluorescence (XRF) spectromicroscopy technique used to perform a comparative analysis of sodium containment in intracellular and interstitial spaces of cardiac tissues taken from animals selected by low and high sodium intake rates. The experimental results obtained show that high sodium daily intake can result in a remarkable increase of sodium concentration in the myocardial interstitium.

## Introduction

The sodium homeostasis processes are traditionally the focus of various studies and experiments. Thus, there is a large amount of evidence showing the negative impact of sodium overloading on myocardia function in the form of its lowered diastolic function [1]. However, recent studies indicate that homeostasis of sodium is much more complex than previously thought: it includes not only osmotic intracellular and extracellular compartments but also a compartment of the interstitial space in which bound Na+ is stored without commensurate water accumulation [2-4] (“three-compartment model”).

As early as in the early 20^th^ century it has been hypothesized that negatively charged glycosaminoglycans (GAGs) of the extracellular matrix may bind to positively charged cations [5, 6]. Thus, sodium becomes osmotically neutral and does not affect the values of arterial pressure (AP) and extracellular fluid volume. We should mention also a process was found to involve the macrophages in response to excessive concentration of sodium in GAG [7].

In 2003 the first experimental proof of water independent sodium storage was delivered by J.Titze *et al*. for rat skins [3, 4]. In 2007-2011 in the framework of the MARS-500 program [8], a prolonged and accurate study on human sodium intake/excretion ratio has found out that amounts of excreted sodium are not related directly to the sodium intake and arterial pressure values. This fact was explained by the existence of osmotically passive sodium deposits in skin and muscle tissues that have been formed earlier under conditions of high sodium diet and made their contribution to the elevated level of sodium in the urine.

To confirm an occurrence of sodium depositions inside striated muscles C. Kopp *et al*. [9, 10] and M. Hammon *et al*. [11] used ^23^Na Magnetic Resonance Imaging (MRI) for studying skin and muscles of the lower legs. Additionally, several MRI studies leading to the conclusion, that the heart is a part of the body’s sodium storage (see, for example, [12]) and showing a link between sodium intake and left ventricular hypertrophy [13]. (Christa et al 2019, European Heart Journal Cardiovascular Imaging).

Our previous investigations with the help of acoustical and X-ray fluorescence analysis (BRUKER S8 Tiger 1 kW WDXRF spectrometer) revealed a correlation between elevated sodium concentration in heart muscles and arterial hypertension as well as patient’s age [14]. We found out also evidence of sodium-related modification of muscle elasticity of myocardia and related diastolic function.

However, all the previous investigations on sodium accumulation in muscle tissues made with the help of MRI or XRF spectrometers were not able to localize the elevated sodium concentration with an adequate high spatial resolution at the cell level. None of the above-mentioned studies could differentiate between extra and intracellular sodium.

The aim of the presented paper was to detect and to estimate the relative concentration of sodium depositions in the extracellular matrix of myocardial tissues of rats with the reference to their sodium consumption rates.

## Results

The samples of cardiac muscle tissues of rats were studied by means of 3 techniques: histology analysis, synchrotron radiation based X-ray transmission microscopy (STXM) and X-ray fluorescent microscopy (XFM) using the same spatial resolution of about 1 micron.

The histological analysis of the tissues taken from high sodium diet animals revealed several interstitial lymphocytes, degranulated mast cells, macrophages, fibroblasts as well as perivascular edema of the interstitium (see Figure1). In contrast to that, no macrophages were found in the tissues of the control group of animals, while the number of fibroblasts was being reduced and mast cells showed no signs of degranulation.

**Figure 1.**
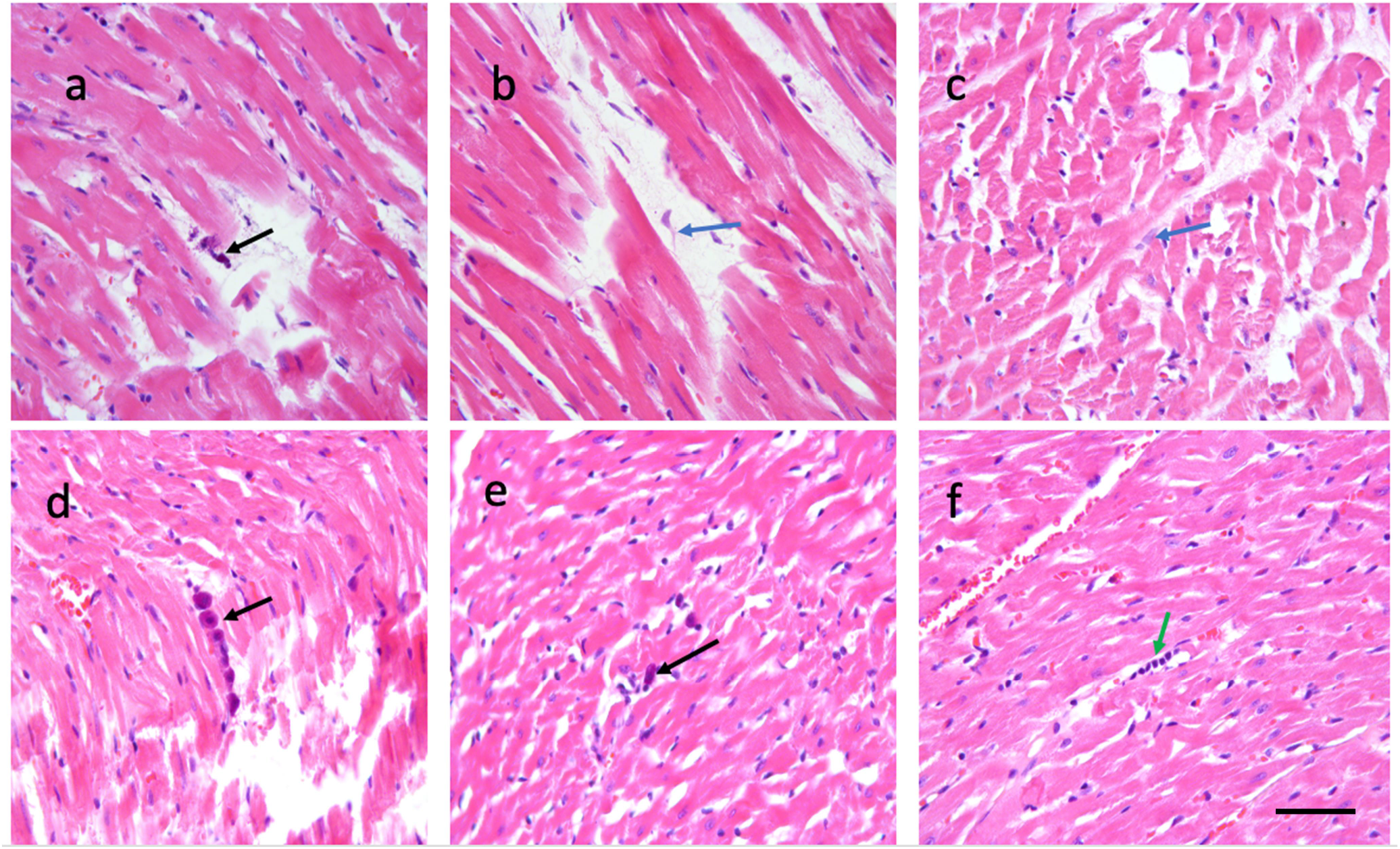
**Optical microscopy images of anomalies in heart tissues of rats from high sodium diet group (images *a* – *c*) in comparison with tissues of control animals (images *d – f*). Black arrow in image *a* shows a degranulated plasmocyte, while blue arrows in images *b* and *c* mark macrophages in the interstitium. In images *d* and *e* one can see also non-granulated mast cells (black arrows). Lymphocytes are marked in image *f* by green arrow. All sections are stained by hematoxylin and eosin. The scale bar for all the images is 150 micron.**

Figures 2 and 3 show typical X-ray absorption images and corresponding sodium maps acquired with the help of synchrotron based STXM and XRF microscopy for heart tissue sections of respectively high and low sodium diet animals. In total 21 ROIs on 7 samples of rat myocardial tissues have been studied: 5 samples were extracted from 5 animals of the high sodium diet group and 2 were taken from 2 animals of the control group. The obtained STXM and XRF microscopy images enabled a clear visual segmentation of cardiomyocytes and intercellular spaces in the heart muscle tissues. In all samples, the XRF analysis revealed a higher concentration of the sodium content inside the cells than in the interstitium space.

**Figure 2.**
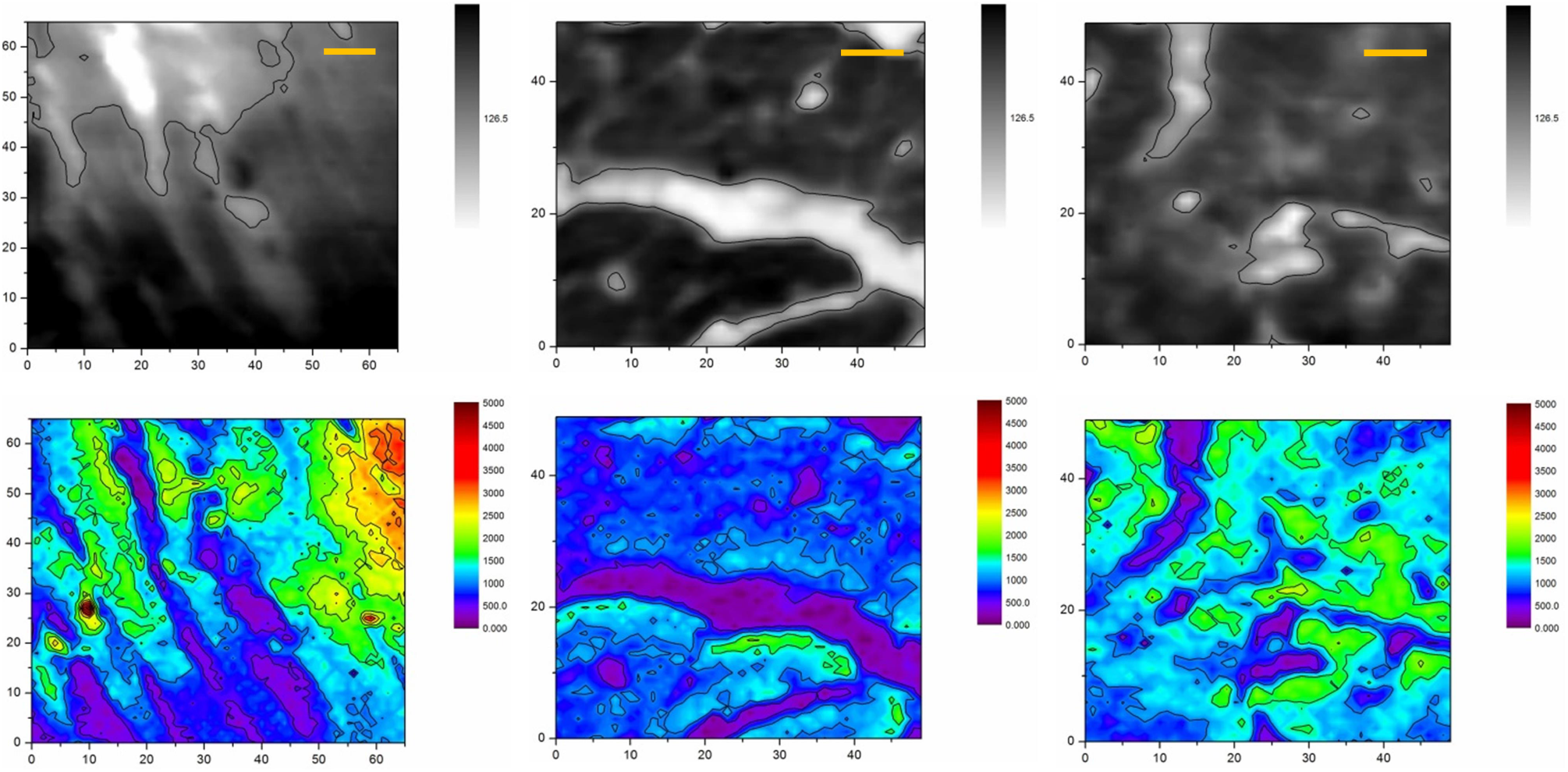
**X-ray images (top row) and sodium XRF maps (bottom row) of heart muscles taken from animals of low sodium diet (control) group. Dark zones (high X-ray absorption) and light zones (low X-ray absorption) correspond to intercellular and extracellular spaces, correspondingly. The scale bars are 10 micron.**

**Figure 3.**
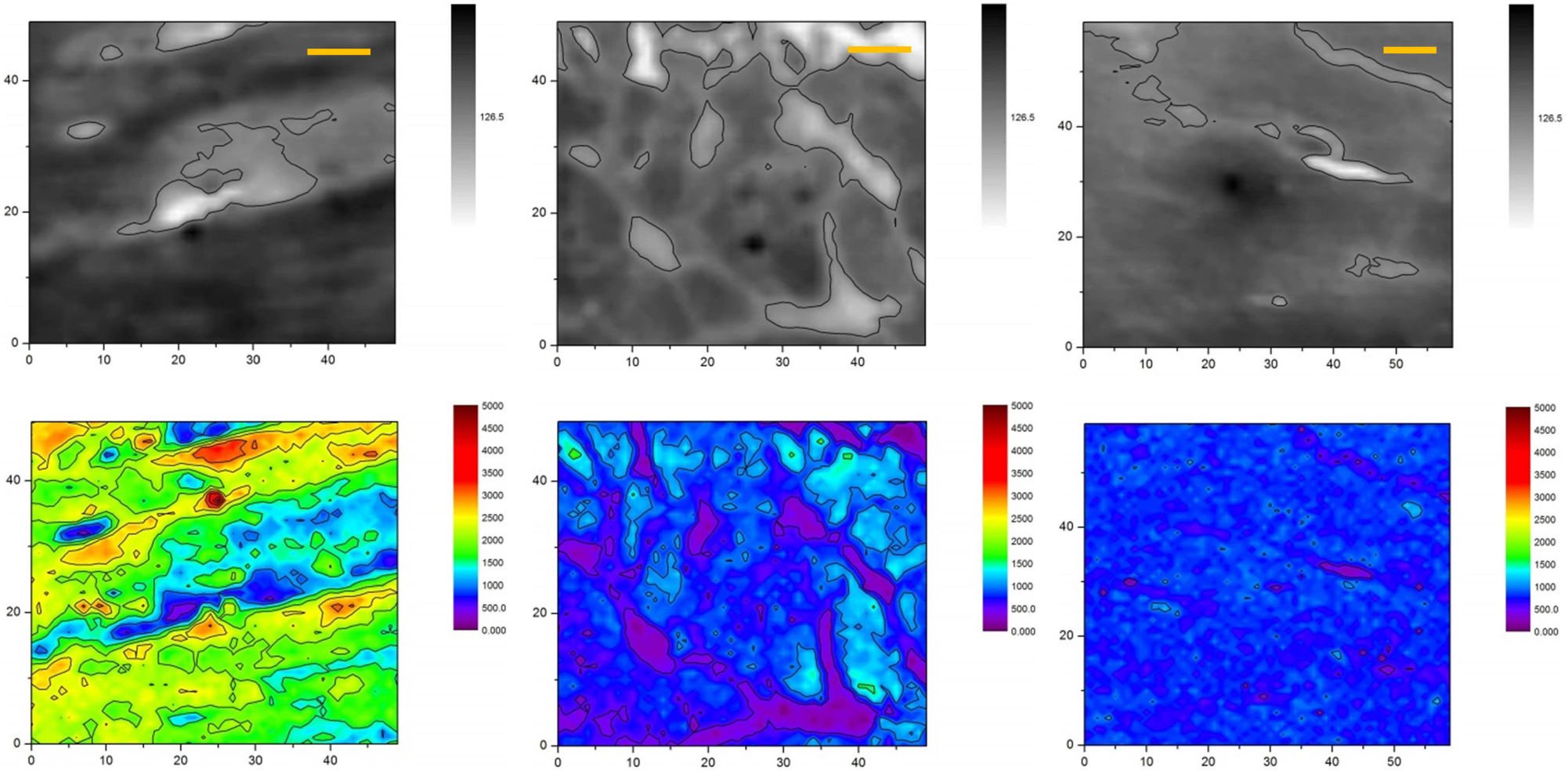
**X-ray images (top row) and sodium XRF maps (bottom row) of heart muscles taken from animals of high sodium diet group. Dark zones (high X-ray absorption) and light zones (low X-ray absorption) correspond to intercellular and extracellular spaces, correspondingly. The scale bars are 10 micron.**

This inverted correlation between intracellular and extracellular sodium can be explained by a known change of light metals content during the chemical fixation of samples. Thus, a side-by-side comparison of XRF elemental maps of cells subjected to different dehydration and drying procedures has demonstrated a significantly reduced amount of highly diffusible elements like *K, Br, Na*, and *Cl* after aldehyde-based conventional fixation [15-17]. A similar disruption of their chemical integrity in chemically fixed or poorly cryofixed cells was observed in secondary ion mass spectrometry (SIMS) experiments [18]. It should be noted that the mentioned problem has been successfully resolved by using methods of fast cryo-fixation.

One can expect that osmotically active fluid sodium would be removed from interstitium and, to a lesser extent, from intracellular volumes restricted by cell membranes. However, in our particular experiments, a wash-out of fluid sodium during the chemical fixation procedure plays a positive role: it reduces significantly a high background level of osmotic sodium of extracellular fluids to express tracks of bound sodium deposits. Due to the comparative character of the presented study and identical treatment of all the samples we do not need the absolute value of local sodium concentration and its extrapolation to in-vivo one.

One can see that sodium is accumulated mostly inside the cells with low traces in interstitium but the extracellular spaces of high sodium diet animals are filled with noticeably more sodium. To estimate this effect quantitatively we applied Kruskal–Wallis test based statistical analysis to the data obtained by XRF microscopy. The main challenge in the statistical processing was to perform an accurate data sampling inside and outside boundaries of cardiomyocytes in the context of thin slice 3D-to-2D transformation of muscle fibers and possible interstitium perforation. This problem has been solved by manual sampling of six 3×3 pixels (3.6×3.6 µm^2^) zones in every ROI to present 3 values of intracellular and 3 values of extracellular sodium content (see Figure 4), thus forming a data sample of sodium K-line emission counts and their averaged intracellular and extracellular values for the particular ROI (see Figure 4 and Table 1).

**Table 1.** **Averaged sodium XRF counts measured in 3 intracellular and 3 extracellular points inside ROIs. B1, B2, and B4 mark the tissue samples extracted from high sodium diet animals, K5 and K7 marks the samples extracted from the control group (low sodium diet)**.

|  | Sample | ROI # | intracellular Na |  |  | extracellular Na |  |  |
| --- | --- | --- | --- | --- | --- | --- | --- | --- |
| High sodium diet | B1 | 1 | 742 | 834 | 834 | 1115 | 932 | 1076 |
|  |  | 2 | 438 | 737 | 1097 | 1736 | 1589 | 1611 |
|  |  | 3 | 1330 | 1700 | 1517 | 1823 | 2256 | 2139 |
|  | B2 | 1 | 304 | 367 | 408 | 574 | 629 | 624 |
|  |  | 2 | 945 | 361 | 517 | 758 | 823 | 915 |
|  | B4 | 1 | 951 | 645 | 1041 | 1034 | 1164 | 1015 |
|  |  | 2 | 577 | 413 | 203 | 1058 | 980 | 967 |
|  |  | 3 | 602 | 193 | 476 | 969 | 857 | 603 |
| Low sodium diet | K5 | 1 | 306 | 434 | 527 | 466 | 717 | 498 |
|  |  | 2 | 910 | 501 | 975 | 1579 | 1469 | 1460 |
|  | K7 | 1 | 386 | 1083 | 649 | 917 | 1344 | 697 |
|  |  | 2 | 261 | 238 | 194 | 1084 | 836 | 962 |
|  |  | 3 | 302 | 295 | 387 | 668 | 694 | 540 |

**Figure 4.**
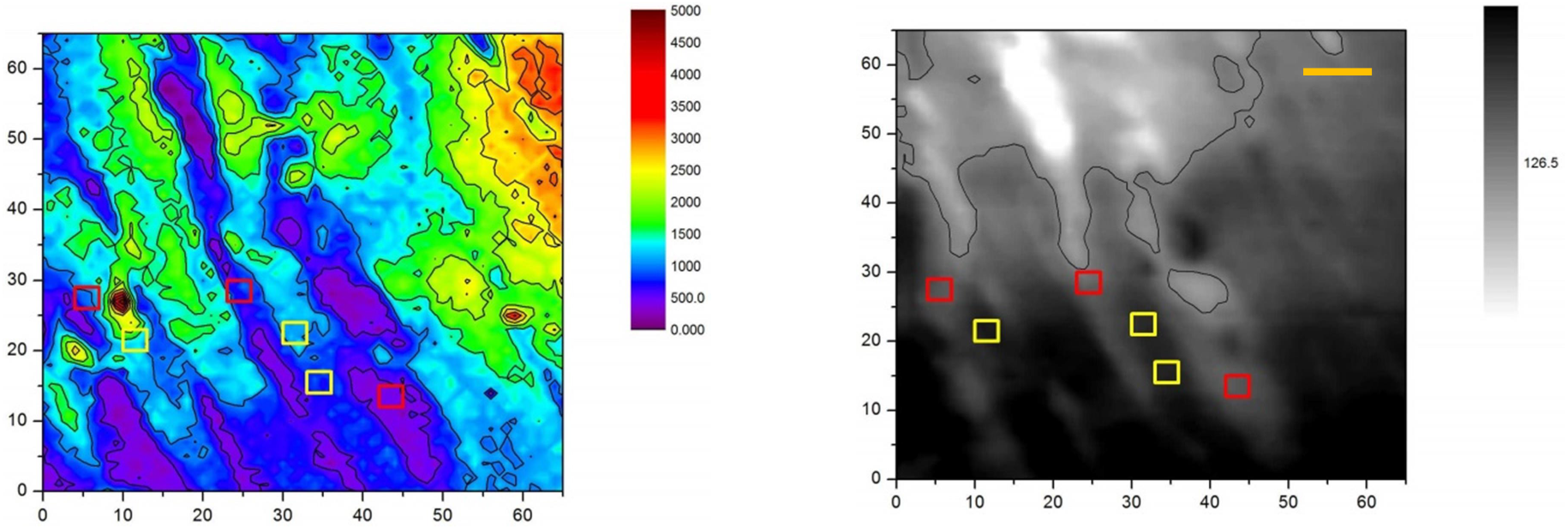
**Illustration of the data sampling procedure based on combined XRF (left) and SXTM (right) images: 3×3 pixels zones were selected in extracellular (red squares) and intercellular (yellow squares) spaces. The scale bar is 10 micron.**

As mentioned above, due to the chemical treatment used for the sample preparation, the amount of sodium inside the cells of all the samples and animals was found to be always higher than in interstitium: 1055.7 ± 445.4 against 632.8 ± 373.6 (mean value, p = 0.000012).

One can suggest that in the case of the absence of sodium deposition bound to GAG structures in the interstitium we should detect nearly the same amount of residual sodium traces in all tissues, which is defined by homeostasis control and identical sample preparation and treatment method used. However, the averaged content of sodium measured in heart tissues for animals fed with high sodium intake rate exceeds that of control animals: 926.6 ± 483.4 against 712.6 ± 394.0 (mean values, p = 0.044).

The results of the statistical processing on the measured data are summarized in Figure 5 where we present a box plot of the sodium contents measured in intracellular and extracellular spaces of heart tissues taken from different groups of animals. One can see no significant difference in the amounts of sodium content inside the cells of both high sodium and low sodium diet groups: 1135.2 ± 474.0 and 928.6 ± 375.7, correspondingly (mean values, *p* = 0.13). However, increased salt intakes result in a clearly elevated level of sodium content left in interstitium outside cardiomyocytes: 718.0 ± 402.4 for high sodium diet in comparison of 496.6 ± 283.5 for low sodium diet (mean values. *p* = 0.037, one low reliable outlier discarded).

**Figure 5.**
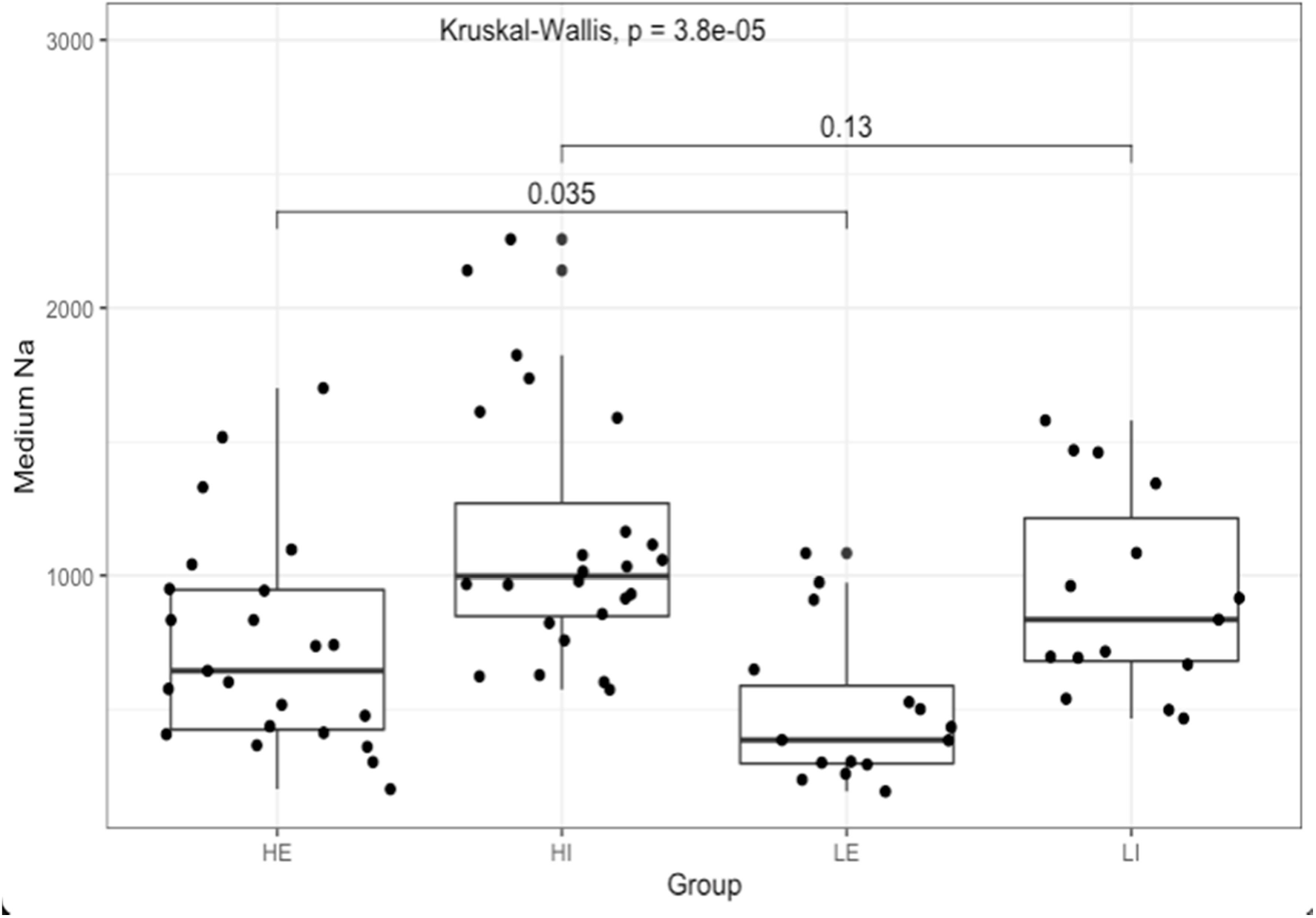
**Box plot of sodium content measured in intracellular and extracellular spaces of heart tissues taken from animals fed with different rates of sodium intake. HE (High Extracellular) and HI (High Intracellular) designate data groups obtained for extracellular and intracellular contents of sodium of high sodium intake diet rats, LE (Low Extracellular) and LI (Low Intracellular) of low sodium diet rats.**

## Discussion

Earlier investigators have studied sodium stored in tissues of biological models and humans by using flame photometry or MRI, with the MRI visualizing an overall elevated content of the tissue or organ. So far, no high spatial resolution imaging of the sodium accumulations at the cell level has been provided. Our study was aimed at more accurate localization of sodium stores in interstitium spaces of myocardial tissues of rats fed with a high rate salt diet without additional hormonal stimulation of the sodium intake.

The present investigation appeared to be successful owing to the high spectral and spatial resolution of synchrotron-based XRF microscopy. Thanks to the imaging capabilities of TwinMic microscope, an increase of sodium deposition was identified inside interstitium spaces of myocardium taken from animals with elevated levels of sodium intake. The main problem here was to recognize full and partial tissue perforations as well as different types of 2D-projections of both cardiomyocytes and interstitium volumes. Of course, the promotion of good quality statistics also needs more strengthening by the processing of more samples and animals. The use of complimentary microtomography or XRF tomography would be profitable as well.

Nevertheless, our findings indicate elevated sodium content inside interstitium of myocardia of high sodium diet animals as compared to animals of the control group with low sodium diet. Sodium washing out of the chemical fixation removed soluble sodium from extracellular volumes, while more conservative sodium content inside cardiomyocyte membranes served as a good reference point for validation of identical parameters and treatment conditions of heart muscle slices.

In general, our results are the next step in the understanding of the relationship between stores of osmotically inactive sodium accumulated in human myocardial tissues due to excessive consumption of salt and the emergence of such clinical conditions as left ventricular hypertrophy of hypertensive patients.

## Conclusion

The world’s first high resolution X-ray imaging of myocardial cells by means of transmission and fluorescence X-ray microscopy techniques has been demonstrated. The results show the elevated concentration of sodium in the interstitium of myocardia that can be explained by its “three-compartment” depositing and confirm the original assumption on the existence of osmotically inactive sodium allegedly bound with negatively charged glycosaminoglycans inside the interstitium network.

We believe the reported experimental results are a valuable basis to continue the research of clinically important deposition of sodium in the myocardia. The next step can be combined research and visualization of sodium and GAG structures by using a combination of XRF and FTIR-spectromicroscopy methods to validate glycosaminoglycan-sodium correlation.

## Materials and methods

### Sample preparation and animals handling

The experiments with animals were carried out in accordance with the international rules of the Guide for the Care and Use of Laboratory Animals and were approved by the Laboratory Animal Care and Use Commission and Local Ethics Committee at Pirogov Russian National Research Medical University.

Fifteen male Wistar rats aged 15-16 weeks were divided into two experimental groups possessing similar body weights: 297.4± 68.4 g (low sodium diet) and 252.0 ± 67.4 g (high sodium diet). The sodium consumption rate for these two groups was different starting from the third week. A low sodium diet limited the sodium intake by the value of 0.2 mEq per 200 g of body weight per day, high sodium diet ration contained 2.0 mEq or more per 200 g of body weight per day that can be considered excessive salt consumption [19, 20]. The water supply was always ad libitum (25 ml of de-ionized water for 15 g of food) for all the animals, while the normal food ration being 15 g per 200 g body weight per week.

High sodium diet feeding was maintained in the corresponding group of animals during 8 weeks before the myocardial hypertrophy occurred. After 8 weeks the animals were intraperitoneally injected with methohexital at a dose of 100 mg/kg and sacrificed through decapitation.

We abide by all appropriate animal care guidelines including ARRIVE 2.0 guidelines for reporting animal research. Rats were housed in cages with a 12:12 light:dark cycle and had access to food and water ad lib at all times unless indicated otherwise. Every effort was made to ameliorate animal suffering.

### Histological staining and study

The 3-µm thick slices of the paraffin-fixed myocardia were mounted onto slides and dried (T=60 °C) in the air. Paraffin was removed with ethanol and the slices were stained by hematoxyline and eosine in accordance with its standard procedures. The histological studies were performed with using of Leica microscope (Leica Microsystems DM2500).

### X-ray fluorescence microscopy

To investigate the sodium distributions inside and outside myocardial cells we employed X-ray fluorescence (XRF) microanalysis combined with scanning X-ray absorption microscopy (SXRM) at ELETTRA TwinMic beamline [21], which synchrotron ring operated at the electron energy of 2.4 GeV with 160 mA current. The incident X-ray beam energy was 1.472 keV to ensure the best excitation and detection of the Kα lines of Na atoms at E=1.041 keV.

The TwinMic microscope was operated in scanning mode. A pinhole of 75 µm diameter served as the secondary X-ray source, which was focused into 1.2 μm spot by a zone plate. The size of the probing X-ray beam was chosen as a compromise between spatial resolution, duration of the scan run and dimensions of the region of interests (ROI). The X-ray flux through the pinhole was about 3·10^9^ photons/s. Absorption contrast XRM images were produced with the help of a fast readout CCD camera (Andor Technology) [22], while the simultaneous XRF signal was collected by up to eight silicon drift detectors. The exposure time was 5 s and seven detectors were used for XRF registration. Each detector had a solid acceptance angle of 3·10^−2^ [23].

The TwinMic configuration enabled visualization of the myocardial tissues in respect to the corresponding sodium, magnesium and other elements distributions with precise overlapping of both the morphological and chemical data. The image size varied from 50×50 to 72×72 pixels. In total 19 ROIs selected in seven samples (3 ROIs per sample in all samples except two) of myocardial tissues have been studied using the combined XRF/SXRM microscopy. In addition to fluorescence images, the corresponding SXRM images of cardiomyocytes and interstitial space were produced.

The obtained XRF maps of chemical elements were analyzed by using open source PyMCA software [24].

### Statistical Processing Methods

For statistical processing of the data we used R language in the *RStudio* software environment (readxl, psych ggplot2 [25], ggpubr, dplyr and tidyr). The check for normality of distribution was carried out using the Shapiro-Wilk criteria and distribution graphs and qq-plot. The methods of nonparametric and parametric statistics were applied to calculate the quantitative indicators as mean values, standard deviations and medians (25 and 75 percentiles). The differences between the data clusters were checked by the Kruskal-Wallis and Mann-Whitney tests. On testing statistical hypotheses we rejected the null hypothesis with a significance level less than 0.05.

## Acknowledgements

The authors thank Dr. N.L.Popov (LPI RAS) for his help and useful comments regarding the study and manuscript. This work was supported by CalipsoPlus EU network travel grant (20185066). “Tecnopolo di nanotecnologia e fotonica per la medicina di precisione” (funded by MIUR/CNR, CUP B83B17000010001) and the TECNOMED project (funded by Regione Puglia, CUP B84I18000540002) are acknowledged for financial support as well. Part of the research reported in this publication was also supported in the frames of HORIZON 2020 Program.

## Author contributions

G.A., D.D., and A.S. devised the main conceptual ideas of the study. D.D., A.S., M.B., and V.M. contributed to sample preparation. A.G, I.A., I.B., M.F., and R.F. conceived, planned and carried out the XRF experiments. G.A., D.D., A.S., A.C., M.F., I.A., and A.V. contributed to the analysis and interpretation of the results. I.A., D.D. and A.G. processed the experimental data, drafted the manuscript and designed the figures. All authors discussed the results and contributed to the final manuscript.

